# Effect of Silicon Application on Leaf mineral Composition of Eucalyptus

**DOI:** 10.1101/2021.01.31.429072

**Authors:** Rahul Kumar, Dilip Kumar Reddy, Prashant Kaushik

## Abstract

Eucalyptus is a commonly planted tree in North India and its cultivation area is increasing. Leaves of Eucalyptus are used for tea which has healing properties and many other economic uses. Silicon is the second most abundant element in the earth’s crust next to oxygen. Silicon (Si) can improve photosynthesis, decrease transpiration, increase water use efficiency and provide resistance against biotic and abiotic stress. The clear information was not available regarding Silicon nourishment in tree species like Eucalyptus. Although a positive in plant development and overall dry matter production was noticed in Eucalyptus with Si application. The amount of nitrogen (N), phosphorus (P), and potassium (K) were inclined leaf whereas Calcium (Ca) and magnesium (Mg) content was not affected by Si application potassium silicate (K_2_SiO_3_). Finally results revealed that upon Si application significant increase in the leaf mineral content in almost all genotypes was recorded. Correlation studies revealed that leaf N and P were highly and significantly correlated with leaf copper (Cu) and iron (Fe) content, leaf Mg and leaf Cu were positively and significantly correlated with leaf N, manganese (Mn) and Fe content, leaf Mn and Fe content were highly and significantly correlated with leaf N, Mg and Cu content. To our detailed knowledge this is the first ever study conducted in Eucalyptus. Here the influence of potassium silicate on popular North Indian cultivars’ leaf mineral content Nevertheless, more studies are required to analyze the effect of Si application in tree species like Eucalyptus.

## Introduction

Eucalyptus is an evergreen tree. Although it is indigenous to Australia, because of its inherent diversity has resulted in successful introduction of many of the species to areas within the tropical, subtropical, and warm temperate zones of the world, for landscape, fuelwood, and timber purposes (Davidson, 1979; Kumar *et al*., 2019; Kumar *et al*., 2020). It has a gum infused bark, incredibly long stems, combined with circular leaves that are difficult to digest. Nevertheless, Eucalyptus leaves could be turned directly into a tea that is risk-free for ingestion. Furthermore, the vegetation may be converted into essential oil for topical use or perhaps inhalation. Tea made from the Eucalyptus leaves has healing properties (Khan *et al*., 2020).

Silicon is primarily discovered as Silicon dioxide (silica Silicates and) that have Silicon, Minerals and Oxygen and is absorbed as silicic acid (H_4_SiO_4_). Epstein (1999) claimed that Silicon is a semi-essential component for crops because deficiency causes abnormal development, reproduction, and development. Silicon triggers defence mechanisms against biotic and abiotic stress and, hence, favorably improves the crop yield and quality. Si has valuable consequences on numerous species of plants, under drought, radiation, high temperature, and freezing (Crusciol *et al*., 2009). It is deposited as amorphous silica in epidermal cell wall, lessens the plant’s water usage, makes more erect leaves, and offers higher photosynthetic rates (Campoe *et al*., 2013) and affects transpiration (Whitehead and Beadle, 2004). Si also reduces the fungal and bacterial incidence by forming a physical barrier (Datnoff *et al*., 1997) and activating some defense mechanisms (Ma and Yamaji, 2006).

Several researchers studied the effect of silicon on trees, and reported an increase in photosynthesis, stimulating root development, improves WUE. Nevertheless, almost all soil-grown crops have Si (Epstein 1994). The uptake depends upon soluble Si of the earth, coming from pedogenic and lithogenic silicates. The weathering procedure’s program is coming from amorphous phytogenic silicates produced from organic and natural content through decomposition processes. Amorphous silica elements, known as phytoliths may be deposited anyplace of the vegetable, within or perhaps between cells and together with the cell wall structure can’t be moved following deposition (Piperno 2006; Raven 1983).

The soils of forest plantations usually have few physical limitations, but low fertility, which requires fertilizer application at appropriate rates and periods for better yields. Some major nutrients N, P and K are applied to eucalyptus plantations (Smethurst *et al*., 2003; 2004). Soil-available phosphorus (P) is generally little, especially in acid soils, thanks to its low solubility, slow diffusion, and sorption (Marschner, 1995). Over 50 % of the evolved land across the earth is P deficient (Vance, 2001). Although changes in foliar mineral concentrations are proficient at mirroring efflux and the influx of nutrition.

Therefore, we hypothesized that senescence could be caused by artificial shading of leaves, that synthetic shading might be utilized to study translocation within senescing vegetation. Synthetic shading has been utilized to bring about translocation additionally to senescence of Cu from leaves of wheat (Hill *et al*. 1979; Teshome *et al*., 2020). There’s a lack of information regarding Si nourishment in tree species like Eucalyptus. Although a positive in plant development and overall dry matter production was noticed in Eucalyptus with Si application. Nevertheless, more studies are required to analyze the effect of Si application in tree species like Eucalyptus. This study is mainly emphasized on the effect of Si application on the leaf mineral content of *Eucalyptus tereticornis*.

## Material and Methods

### Study Area and Plant material

Study was performed under a greenhouse located at Agriculture Research Farm, Kurukshetra, Haryana, India coordinates at 29.94°N 76.89°E the seedling of 5 Eucalyptus varieties were obtained from the Haryana Forest Department and 2 Genotypes were procured from the WIMCO Ltd, Uttarakhand, India (Table 1). There were in total 3 replications of individual genotypes comprising 5 plants in each replication. After acclimatization, the seedlings of individual variety were selected based on the height and collar diameter and were transferred to individual plastic pots (3 L), with a mixture of coco peat vermiculite and perlite. Treatments were applied for 60 days. The treatments were arranged in a randomized complete block design. The silicon application as potassium silicate (K_2_SiO_3_), at the dosage of 1.00 mmol Si L^−1^ was compared against control.

**Table 1.** Eucalyptus genotypes used in the present study.

| Genotype | Species | Source |
| --- | --- | --- |
| BCM 411 | *Eucalyptus tereticornis* | ITC, Bhadrachalam Paper Mills, Telangana, India |
| BCM 2045 | *Eucalyptus tereticornis* | ITC, Bhadrachalam Paper Mills, Telangana, India |
| BCM 413 | *Eucalyptus tereticornis* | ITC, Bhadrachalam Paper Mills, Telangana, India |
| Wimco 12 | *Eucalyptus tereticornis* | WIMCO Ltd., Uttarakhand, India |
| Wimco 14 | *Eucalyptus tereticornis* | WIMCO Ltd., Uttarakhand, India |

### Mineral Content Estimation

The composite samples were ground separately and examined for various nutrients.

### Estimation of N content

Nitrogen (N) content in the leaf tissues of Eucalyptus was estimated using the method proposed by Kjeldahl method (Piper 1950).

### Estimation of P content

Phosphorus (P) was examined by phosphomolybdic blue colorcolorimetric technique by using barton’s reagent and intensity of the colour was measured at 420nm in spectrophotometer (Jackson, 1958).

### Estimation of K, Ca, Mg, Cu, Fe, Mn, and Zn

Potassium (K), calcium (Ca), magnesium (Mg), copper (Cu), iron (Fe), manganese (Mn) and zinc (Zn) - were estimated with the help of atomic absorption spectrophotometry in extracts received through digestion of 0.5 g of dried leaf sample within a potent acid blend HNO_3_/H_2_SO_4_/perchloric acid (Nikalus *et al*., 2006; Wisdom *et al*., 2016).

### Statistical Analysis

Means of all treatments, i.e., with and without Si application were subjected to analysis of variance (ANOVA) to estimate the differences using SPSS software (11.5 version). The significant differences among treatment means were then assessed by Duncan’s Multiple Range Test (DMRT) for comparison of variance separated with least significant difference (LSD) as a post hoc test. Pearson’s correlation coefficients were determined among the mineral grand mean using the JASP (version 0.14.1) and the network analysis was also performed using JASP via EBIC glasso estimator with a bootstrap of 1000 to determine the strength of relation among the minerals.

## Results

### N, P, K Content

Under the influence of Si, the Eucalyptus leaves showed almost 100% more accumulation of N (Figure 1). The top three genotypes BCM 2045, BCM 411 and BCM 413 followed by the WIMCO series genotypes (Figure 1). P content with the application of silicon was improved, whereas the genotypes BCM 413 and WIMCO 12 were least affected by the silicon application (Figure 1). BCM 413, WIMCO 12 and WIMCO 14 were the accessions with the highest P content (Figure 1). The leaf P content was maximum in the treatment of BCM 413 and WIMCO 12 (Figure 1). In contrast, the topmost responses to Si application were BCM 413 and WIMCO 14 (Figure 1). Genotypes with the highest amount of leaf K concentration were BCM 2045, BCM 411 and WIMCO 14. Simultaneously, the most responsive accessions to the Si application were BCM 413 and WIMCO 12 (Figure 1).

**Figure 1.**
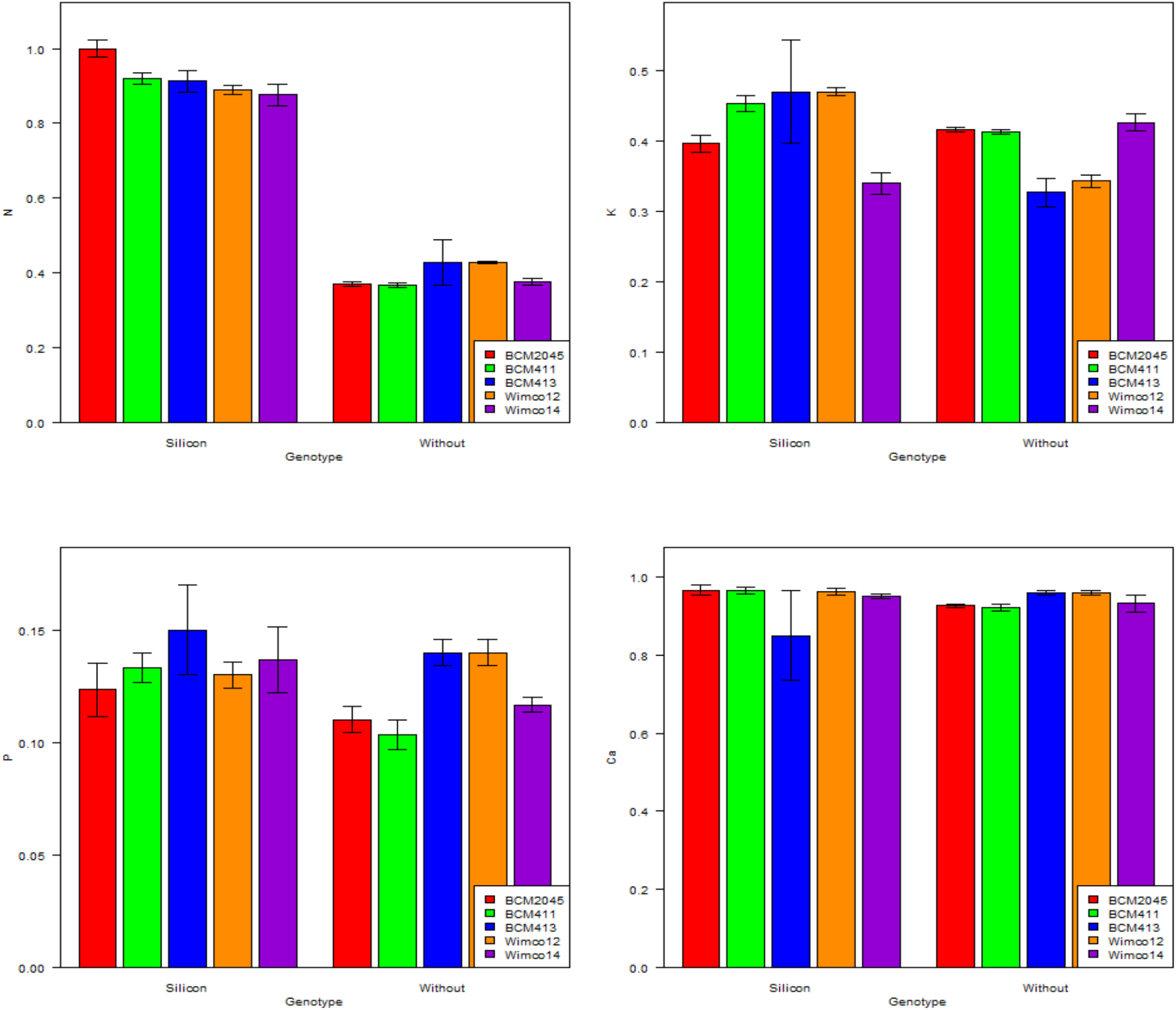
Variation determined for the N, P, K and Ca content of Eucalyptus leaves with and without Si application.

### Ca, Mg, Cu, Fe, Mn, and Zn Content

Surprisingly there were no significant differences for the Ca content as if it remained the same except for the genotype BCM 413 (Figure 2). Moreover, the leaf Mg content concentration remained mostly unaffected by the Silicon application except for all the accession with small and similar increments (Figure 2). Almost equal concentration of leaf Cu content was determined in the BCM 2045 and BCM 411. At the same time, the most responsive to Si application was WIMCO 12 (Figure 2). Almost equal concentration of leaf Zn content was determined in the BCM 2045 and BCM 411. In comparison, the most responsive to Si application was WIMCO 12 (Figure 2). All the accessions showed almost equal Zn concentration in the leaves under normal conditions. In contrast, there was a significant improvement for the leaf Mn content with the application of Si. The Mn content in the leaf of accession BCM 411 was almost doubled after the application of Si (Figure 2).

**Figure 2.**
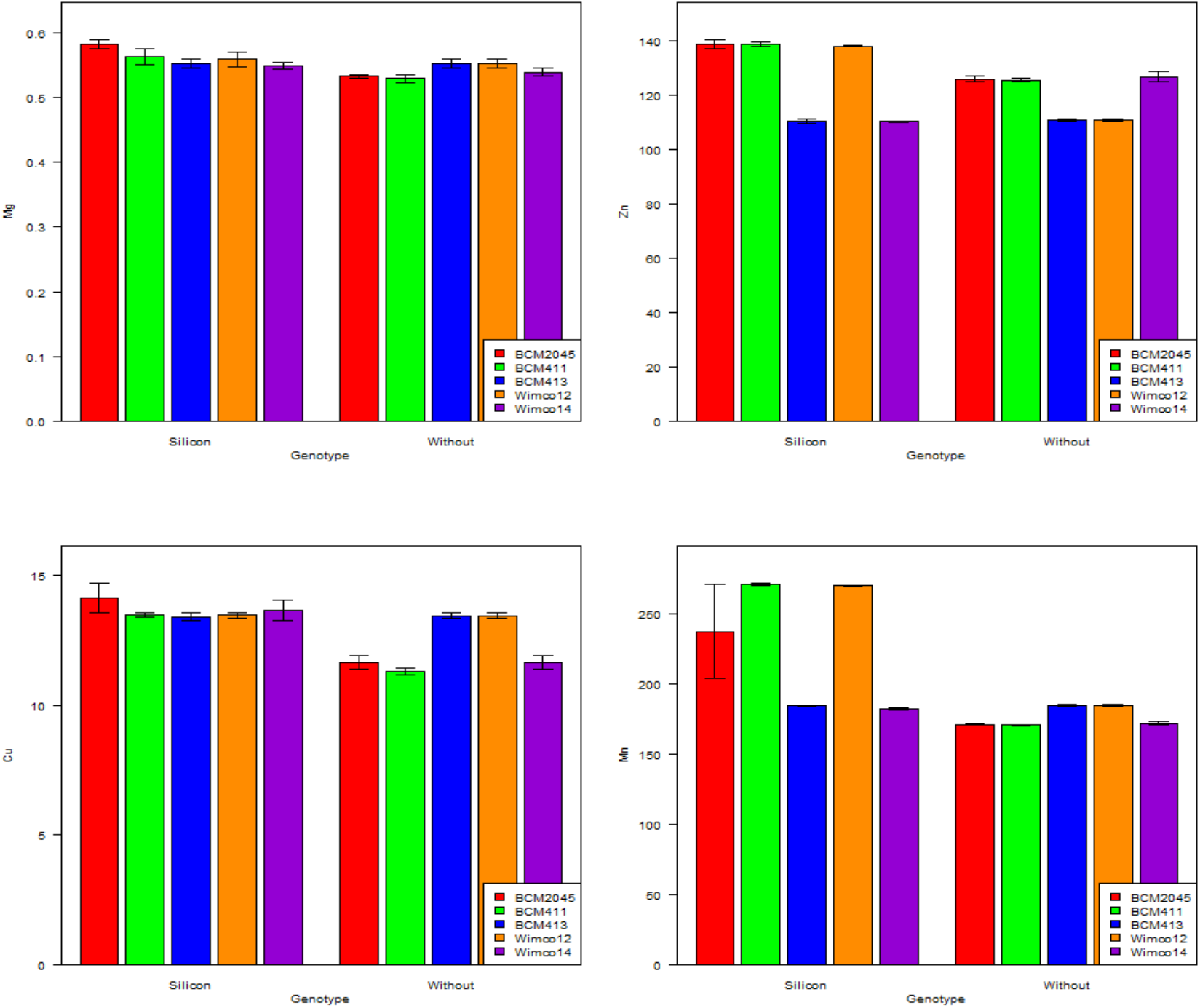
Variation determined for the Mg, Cu, Zn and Mn content of Eucalyptus leaves with and without Si application.

The highest content of leaf Fe content was recorded for the genotypes BCM 413 and WIMCO 12 (Figure 3). Furthermore, their leaf Fe concentration remained unchanged even after the Silicon application (Figure 3). In contrast, BCM 2045, BCM 411 and WIMCO 14 were determined to be very responsive to Si application as their leaf Fe content increased almost 300 % after Si application (Figure 3).

**Figure 3.**
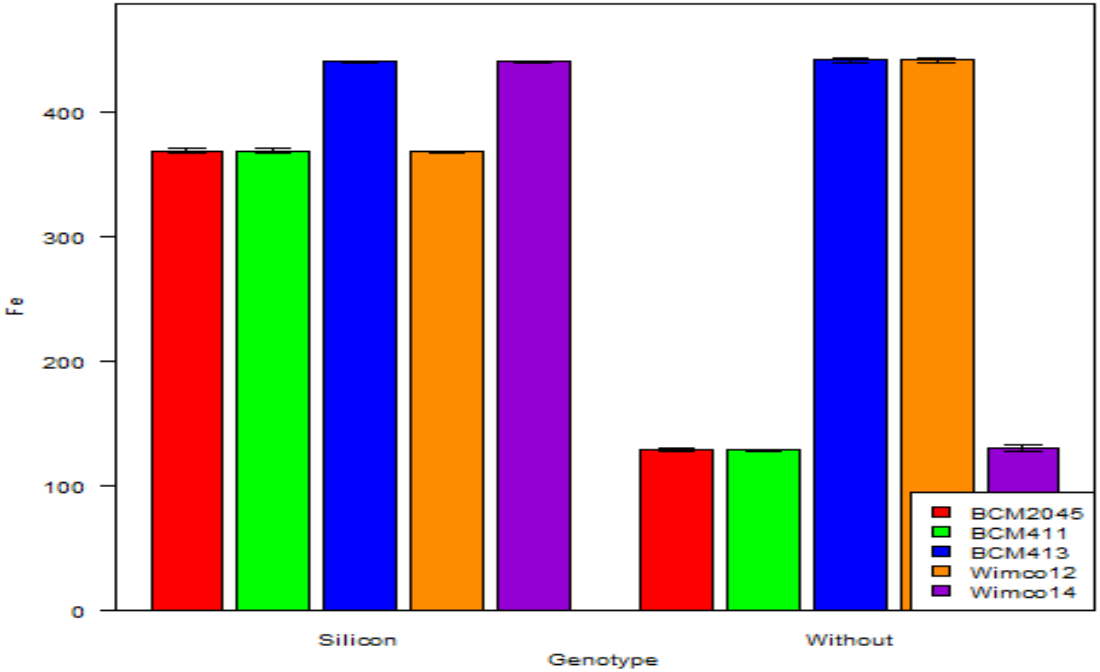
Variation determined for the Fe content of Eucalyptus leaves with and without Si application.

**Table 2.** Variation parameters for the leaf mineral content in Eucalyptus (Figure 3).

|  | Replication | Genotype | Treatment | Genotype × Treatment | Error |
| --- | --- | --- | --- | --- | --- |
| Df | 2 | 4 | 1 | 4 | 18 |
| N (%) | 0.03 | 0.31 | 2.08 | 0.66 | 0.02 |
| F-ratio | 0.64 | 1.54 | 1034.04 | 3.3 |  |
| p-value | 0.5374 | 0.2339 | <0.001 | 0.034 |  |
| P (%) | 0.01 | 0.08 | 0.0012 | 0.0003 | 0.03 |
| F-ratio | 0.46 | 2.68 | 3.84 | 1.05 |  |
| p-value | 0.6404 | 0.0648 | 0.0658 | 0.4109 |  |
| K (%) | 0.19 | 0.02 | 0.12 | 0.14 | 0.02 |
| F-ratio | 0.96 | 0.99 | 6.2 | 7.09 |  |
| p-value | 0.4012 | 0.4383 | 0.0228 | 0.0013 |  |
| Ca (mg/g) | 0.07 | 0.26 | 0.01 | 0.0059 | 0.04 |
| F-ratio | 1.76 | 0.67 | 0 | 1.5 |  |
| p-value | 0.196 | 0.622 | 0.954 | 0.244 |  |
| Mg (mg/g) | 0.01 | 0.02 | 0.03 | 0.07 | 0.02 |
| F-ratio | 0.22 | 1.15 | 16.27 | 3.57 |  |
| p-value | 0.807 | 0.366 | <0.001 | 0.026 |  |
| Cu (mg/g) | 0.1661 | 13.225 | 133.467 | 2.306 | 0.21 |
| F-ratio | 0.79 | 6.32 | 63.77 | 11.02 |  |
| p-value | 0.467 | 0.002 | <0.001 | <0.001 |  |
| Zn (mg/g) | 4.24 | 520.63 | 389.37 | 408.31 | 2.73 |
| F-ratio | 1.56 | 190.61 | 142.55 | 149.48 |  |
| p-value | 0.238 | <0.001 | <0.001 | <0.001 |  |
| Mn (mg/g) | 411.11 | 2895.58 | 20578.53 | 3045.77 | 337.31 |
| F-ratio | 1.22 | 8.58 | 61.01 | 9.03 |  |
| p-value | 0.318 | <0.001 | <0.001 | <0.001 |  |
| Fe (mg/g) | 45.16 | 49448.96 | 15327.42 | 42810.41 | 2.61 |
| F-ratio | 17.35 | 18994.42 | 58861.8 | 16444.4 |  |
| p-value | <0.001 | <0.001 | <0.001 | <0.001 |  |

### Correlation and Network Analysis

Out of the total 36 correlations determined for the leaf mineral content in this study, 13 were significant (Figure 4). Among them 3 were in the negative direction and the remaining 10 were in a positive direction (Figure 4). Leaf N content was positively correlated with the leaf Fe, Mn, Cu and Mg (Figure 4). Whereas leaf P content was negatively correlated with the leaf Ca content (Figure 4). In comparison, a significant and positive correlation was determined between the P, Fe and Cu content of leaf with that of leaf P content. Moreover, the K content in the leaves of Eucalyptus was positively correlated with the leaf Zn content (Figure 4). Furthermore, the Mg content was determined to be positively correlated with the Fe, Mn, and Cu (Figure 4). Whereas, the Cu content was highly correlated with the Fe and the Mn content. Moreover, the correlation between Cu and Fe content of the leaf was absolute. In a similar direction, high correlation was detached between the Zn and the Mn content (Figure 4). The network analysis showed a stronger connection between the mineral with the Si application than the connection under control, which was only vital between the Fe, Zn and Mn and to some degree among the leaf N content with Mg and Cu content of leaf (Figure 5).

**Figure 4.**
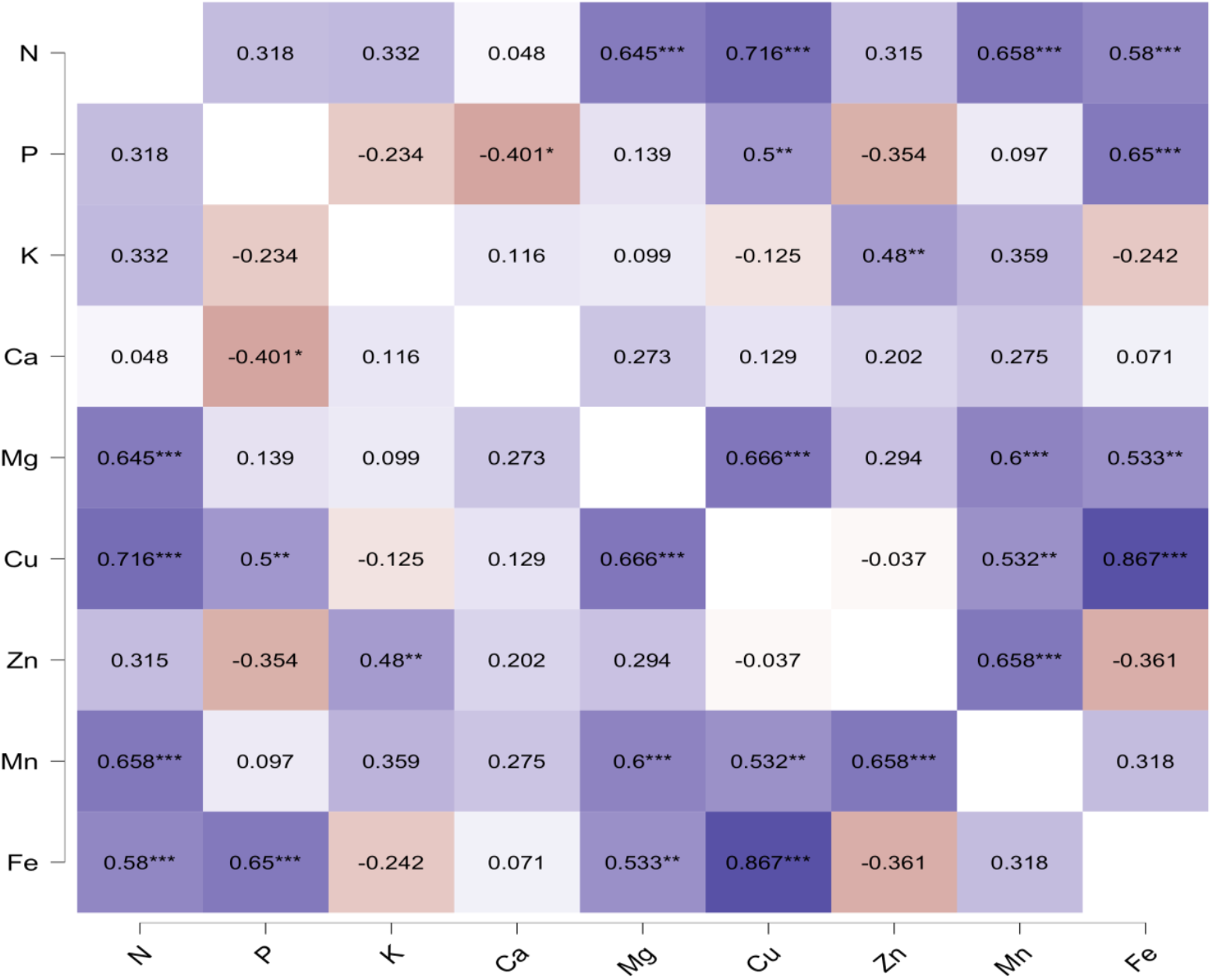
Pearson’s linear correlations coefficient among the mineral content studied in the present study.

**Figure 5.**
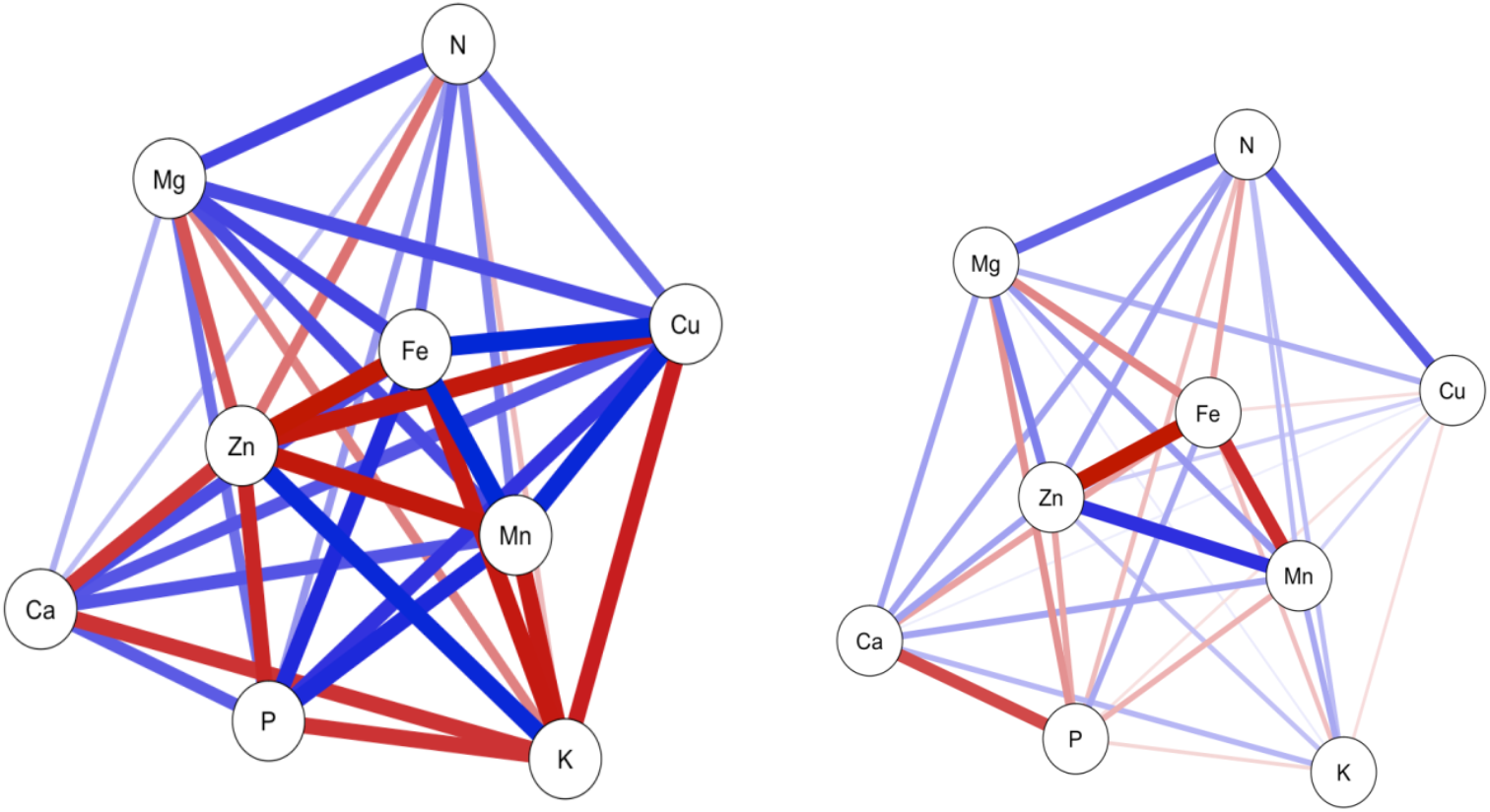
Network analysis under the influence of Si (left) and in the control treatment (right).

## Discussion

Nitrogen content accumulation was more in the leaves in the application of silicon. The N content was maximum in three genotypes BCM 2045, BCM 411 and BCM 413 followed by the WIMCO series genotypes. Similarly, Mehrabanjoubani *et al* (2015) reported that the Si application improved all vegetative and reproductive growth parameters. Silicon application increased Zn as well as Ca^2+^, K^+^, P and B contents in plants. Potassium (K) and phosphorus (P) deficiency decrease physical stability, quality, drought tolerance and crop opposition to pathogens. Potassium deficiency also results in oxidative stress, as established by a buildup of ROS and membrane lipid peroxidation (Cakmak, 2005). P content was improved with the application of silicon, whereas the genotypes BCM 413 and WIMCO 12 were least affected by the silicon application. The leaf P content was found high in the treatment of BCM 413 and WIMCO 12. The beneficial effect of Si on P in rice was clear and showed increased P availability within P deficient plants (Ma and Takahashi, 1990). Nevertheless, the underlying mechanism worried in subsequent exudation and biosynthesis of organic acids in response to P deficiency is not clear (Ryan *et al*., 1997; Kochian et al. 2004; Neumann and Römheld 2007).

Root P uptake is mediated by higher affinity plasma membrane (PM) associated Pi transporters. Beneficial effects of Si under P deficiency has been emphasized for the numerous members of Gramineae family such as wheat. Indeed, the assessments of Takahashi *et al*., 1990; Jianfeng et al., 1991 did not confirm that improving amount of applied Si is able to have an impact on both fixation capacity or possibly accessibility of gardening soil P. Ma (2004) proposed that the larger beneficial effect of Si on plant growing under P deficiency emphasize as a result of the enhanced accessibility of internal P via the drop of excess Fe and Mn uptake. Thus, the leaf P target in Si addressed wheat plants with absolutely no P application achieved the amount of P fertilized vegetation.

Nevertheless, an immediate effect of Si in mediating K deficiency has not been uncovered. The K content was inclined in the Si application. The preliminary exploration of Miao *et al*. (2010) exhibits Si to K deficient soybean (*Glycine max*) vegetation enhanced each internal K status and plant growth. Si additionally relieved K deficiency induced membrane lipid peroxidation in addition to oxidative stress by modulating antioxidant enzymes. The leaf Mg and Ca content concentration remained mostly unaffected by the Silicon application. In contrast, information that’s restricted may be bought on the benefits of Si nourishment under the absence of various other micronutrients and Fe. This is partially an outcome of the stage that root response to Fe deficiency has been examined and mainly characterized in nutrient solution tests where Si was omitted. Recently it has been discovered that the addition of Si on the substance remedy can mitigate Fe deficiency chlorosis in technique one plant life as cucumber, pumpkin.

Nevertheless, Si supply had no adverse effect on zinc’s health problem (Zn) and Mn. However, it decreased leaf necrosis symptoms (Bityutskii *et al*., 2014), which is possibly an outcome of Si’s indirect effect on boosting antioxidant defence ability in plant cells. Application of Si facilitated mobility plus xylem translocation of Fe towards shoot, along with cells buildup of Fe mobilizing elements as citrate (xylem sap, root and shoot tissues) and perhaps catechins (in roots). Reliant on these findings, Si’s treating outcome appears to be a lot more indirect, by negatively affective activation of Fe deficiency associated genes accountable for improved root acquisition and Fe’s cell mobilization (Kaushik and Saini, 2019). Despite, the most plentiful element in the Earth’s crust, metallic (Fe) deficiency is one of the main limiting factors for crop generation in calcareous soils all around the globe (Vose 1982; Romera *et al*., 2019). Although still in progress, these findings are not just brand new proof of the manifold beneficial job of Si in acquiring nourishment. More research is necessary to analyze the effect of Si application in tree species.

## Conclusion

Maximum accumulation of nitrogen was found in the genotypes BCM 2045, BCM 411 and BCM 413 followed by the WIMCO series genotypes under the application of Si. Highest P content was observed in the accessions BCM 413, WIMCO 12 and WIMCO 14. The amount of leaf K was observed increased in Si applied treatments. Highest amount of leaf K concentration was found in BCM 2045, BCM 411 and WIMCO 14. The leaf Ca and Mg content was not affected on Si application. Concentration of leaf Cu and Zn content was almost similar in the BCM 2045 and BCM 411. At the same time, the most responsive to Si application was WIMCO 12 with respect to Cu and Zn. Double increment of leaf Mn content was recorded upon the application of Si in accession BCM 411. The highest leaf Fe content was recorded for the genotypes BCM 413 and WIMCO 12, but silicon application did not affect the Fe content of the leaf. The accessions BCM 2045, BCM 411 and WIMCO 14 were highly responsive to Si application as their leaf Fe content increased almost 300 % after Si application. Correlation study revealed that out of 36 correlations, 13 were significant among these 3 were in the negative direction and the remaining 10 were in a positive direction. Finally, it could be concluded that Si application recorded significant increase in the leaf mineral content in almost all genotypes.

## Notes

### Competing Interest Statement

The authors have declared no competing interest.

